# Tracing the phylogenetic history of the Crl regulon through the *Bacteria* and *Archaea genomes*

**DOI:** 10.1101/410621

**Authors:** A Santos-Zavaleta, E Pérez-Rueda, M Sánchez-Pérez, D A Velázquez-Ramírez, J Collado-Vides

## Abstract

Crl, identified for curli production, is a small transcription factor that stimulates the association of the σ^S^ factor (RpoS) with the RNA polymerase core through direct and specific interactions, increasing the transcription rate of genes during the transition from exponential to stationary phase at low temperatures, and it uses indole as an effector molecule. The lack of a comprehensive collection of information on the Crl regulon makes it difficult to identify a dominant function of Crl and to generate any hypotheses concerning its taxonomical distribution in archaeal and bacterial organisms. In this work, based on a systematic literature review, we identified the first comprehensive dataset of 86 genes under the control of Crl in the bacterium *Escherichia coli* K-12; those genes correspond to 40% of the σ^S^ regulon in this bacterium. Based on an analysis of orthologs in 18 archaeal and 69 bacterial taxonomical divisions and using *E*. *coli* K-12 as a framework, we suggest three main events that resulted in this regulon’s actual form: (i) in a first step, *rpoS*, a gene widely distributed in bacteria and archaea cellular domains, was recruited to regulate genes involved in ancient metabolic processes, such as those associated with glycolysis and the tricarboxylic acid cycle; (ii) in a second step, the regulon recruited those genes involved in metabolic processes, which are mainly taxonomically constrained to *Proteobacteria*, with some secondary losses, such as those genes involved in responses to stress or starvation and cell adhesion, among others; and (iii) in a posterior step, Crl was recruited as a consequence of its emergence in *Enterobacteriaceae*. Therefore, we suggest that the regulon Crl is highly flexible for phenotypic adaptation, probably as consequence of the diverse growth environments associated with all organisms in which members of this regulatory network are present.

## Introduction

Gene expression in bacteria is coordinated through the interplay of sigma (σ) factors on the core RNA polymerase (RNAP) [1] and DNA-binding transcription factors (TFs), providing bacteria with the ability to express multiple genes under different metabolic stimuli or growth conditions. In the bacterium *Escherichia coli* K- 12, seven sigma factors have been experimentally identified, together with around 300 different TFs responsible for recognizing and activating almost all of their genes. Among these, RpoD, or σ^70^, regulates around 40% of the total gene repertoire, whereas alternative sigma factors such as RpoS (σ^S^), the master regulator of the stationary-phase response [2], regulate between 5 and 10% of the total genes in *E*. *coli* K-12 [3,4].

Sigma factors and TFs regulate a large diversity of genes, hierarchically organized in regulons [5]. Previous comparative genomics studies have suggested that regulons exhibit considerable plasticity across the evolution of bacterial species [6]. In this regard, comparison of the gene composition of the PhoPQ regulon in *E*. *coli* and *Salmonella enterica* serovar Typhimurium revealed a very small overlap in both species, suggesting a low similarity in composition between the target genes that are specifically PhoP regulated in *S*. Typhimurium strains and in *E*. *coli* K-12 [7]. Incidentally, this plasticity in bacterial regulons is evidence of lineage-specific modifications [8].

In this regard, we conducted an exhaustive analysis concerning the conservation of the Crl regulon in Bacteria and Archaea cellular domains, using as a reference the currently known system in *E*. *coli* K-12. Contrary to the most common regulatory mechanisms that involve the direct binding to operators or activators, Crl is an RNAP holoenzyme assembly factor that was originally identified in curli production. It is expressed at low temperatures (30°C) [9] during the transition phase between the exponential and stationary phases, under low osmolarity, as well as in stationary phase [10]. In *E*. *coli*, Crl has a global regulatory effect in stationary phase, through σ^S^, as it reorganizes the transcriptional machinery [11], stimulating the association of σ^S^ with the RNAP core, tilting the competition between σ^S^ and σ^70^ during the stationary phase in response to different stress conditions [12, 13] [9, 14]; its production is concomitant with the accumulation of σ^S^ [9].

Assembling the different pieces of the Crl regulon and its regulatory network into one global picture is one of our objectives in this work. The evaluation of this regulon in *Bacteria* and *Archaea* will provide clues about how the regulation of genes by Crl has been recruited in all the organisms, i.e., if the regulated genes were recruited similar to Crl or if they followed different pathways. To this end, 86 genes under the control of Crl in *E*. *coli* K-12 were compiled from exhaustive literature searches. To our knowledge, this is the first attempt to describe the genes regulated by Crl in *E*. *coli* K-12; in addition, Crl homologs were searched among bacterial and archaeal genomes and identified in low copy numbers, constrained to *Enterobacteriaceae* species. Finally, members of the regulon were identified as widely distributed beyond enterobacteria, suggesting that Crl was recruited in a secondary evolutionary event to regulate a specific supset of genes, most likely genes involved in a functional response in enterobacteria to contend against starvation.

## Methods

### Identification of Crl-regulated genes

We performed an exhaustive search of the literature related to Crl in *E*. *coli* K-12 in PubMed [15] under the following search strategy: “coli in the title (to exclude spurious articles) and Crl in all fields,” and “regulation” and “*rpoS*” with different combinations. We obtained 21 manuscripts with relevant information. In addition, genes under the control of Crl were obtained from microarray data analysis with *crl* mutants and with data processed by our authors (Table 1). Finally, we searched for gene/operon notes in RegulonDB and EcoCyc [3, 16] for Crl interactions and σ^S^ promoters; for assembling the network of regulation of Crl; and to identify associations between the TF and regulatory role for each member of the Crl regulon. We must remember that RegulonDB is the main database on transcriptional regulation in *E*. *coli* K-12 of manually curated data from scientific publications.

**Table 1.**
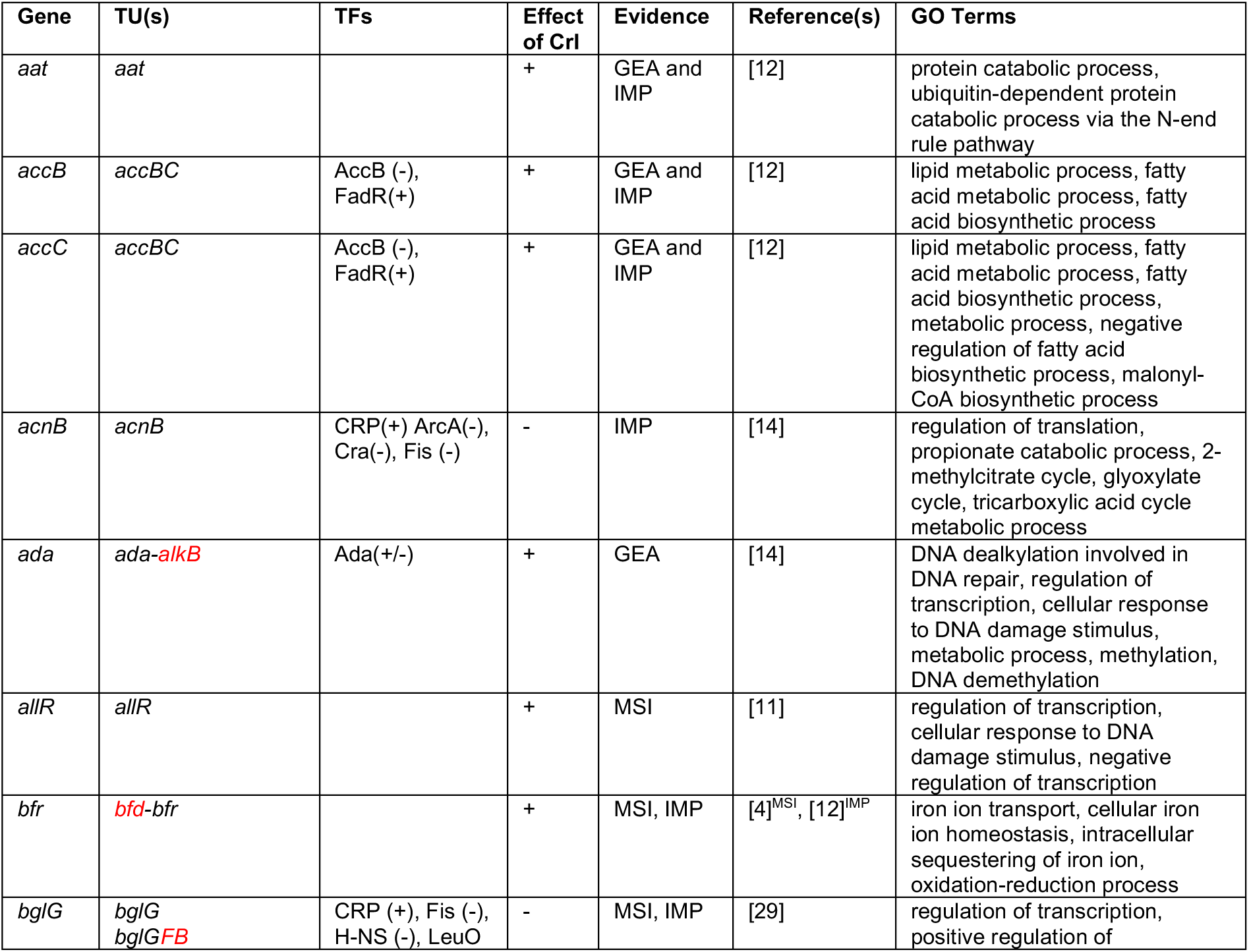
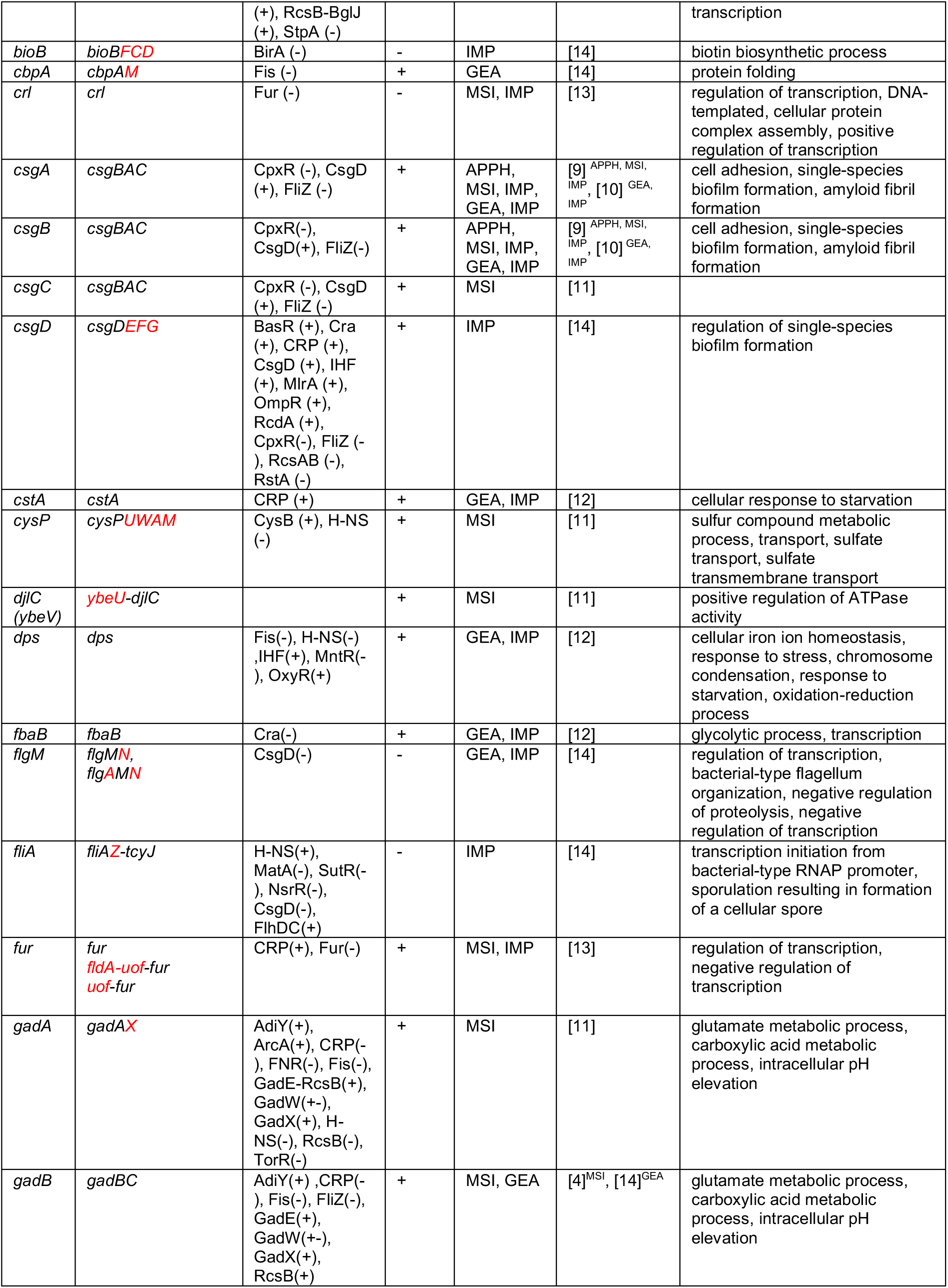
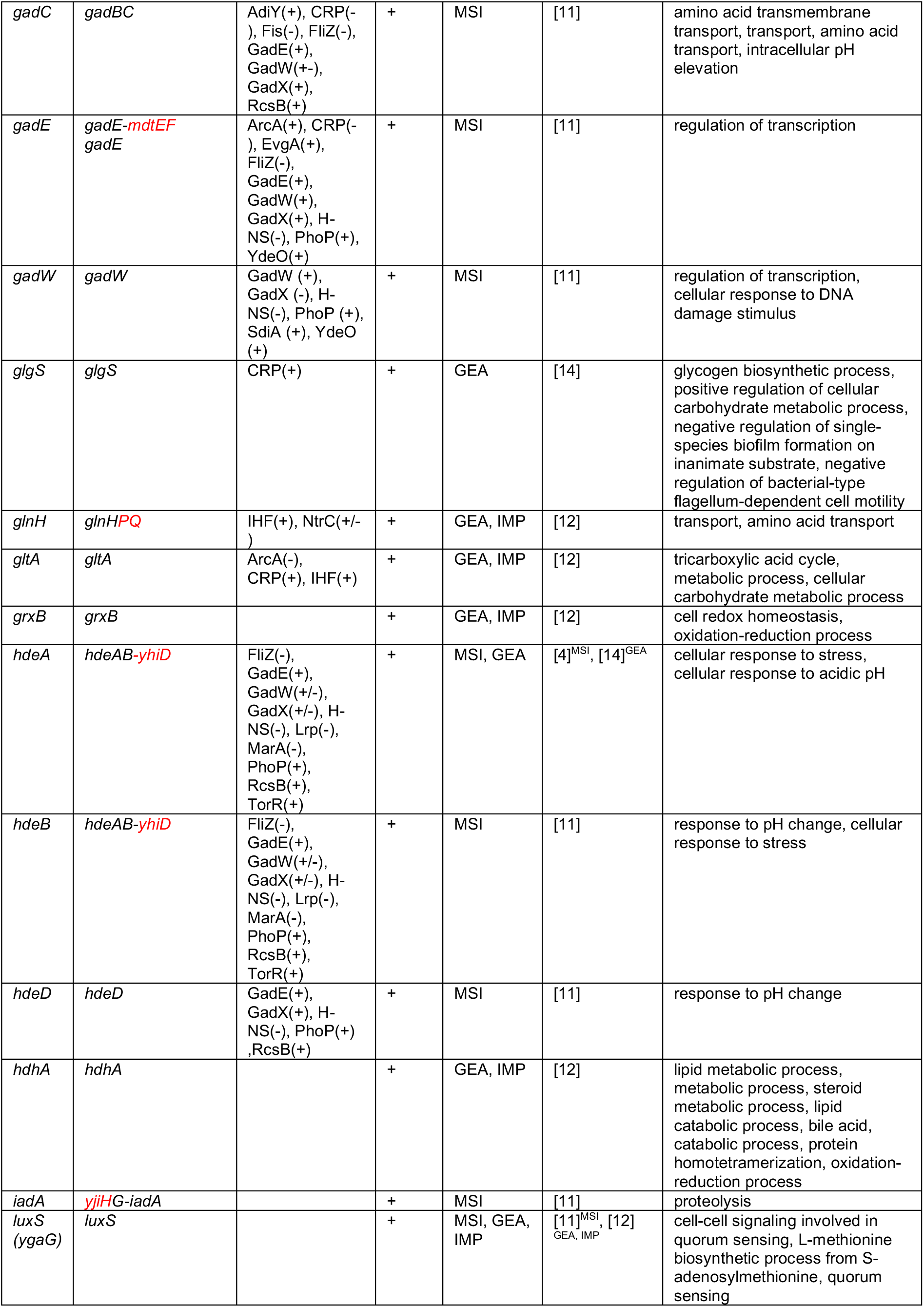
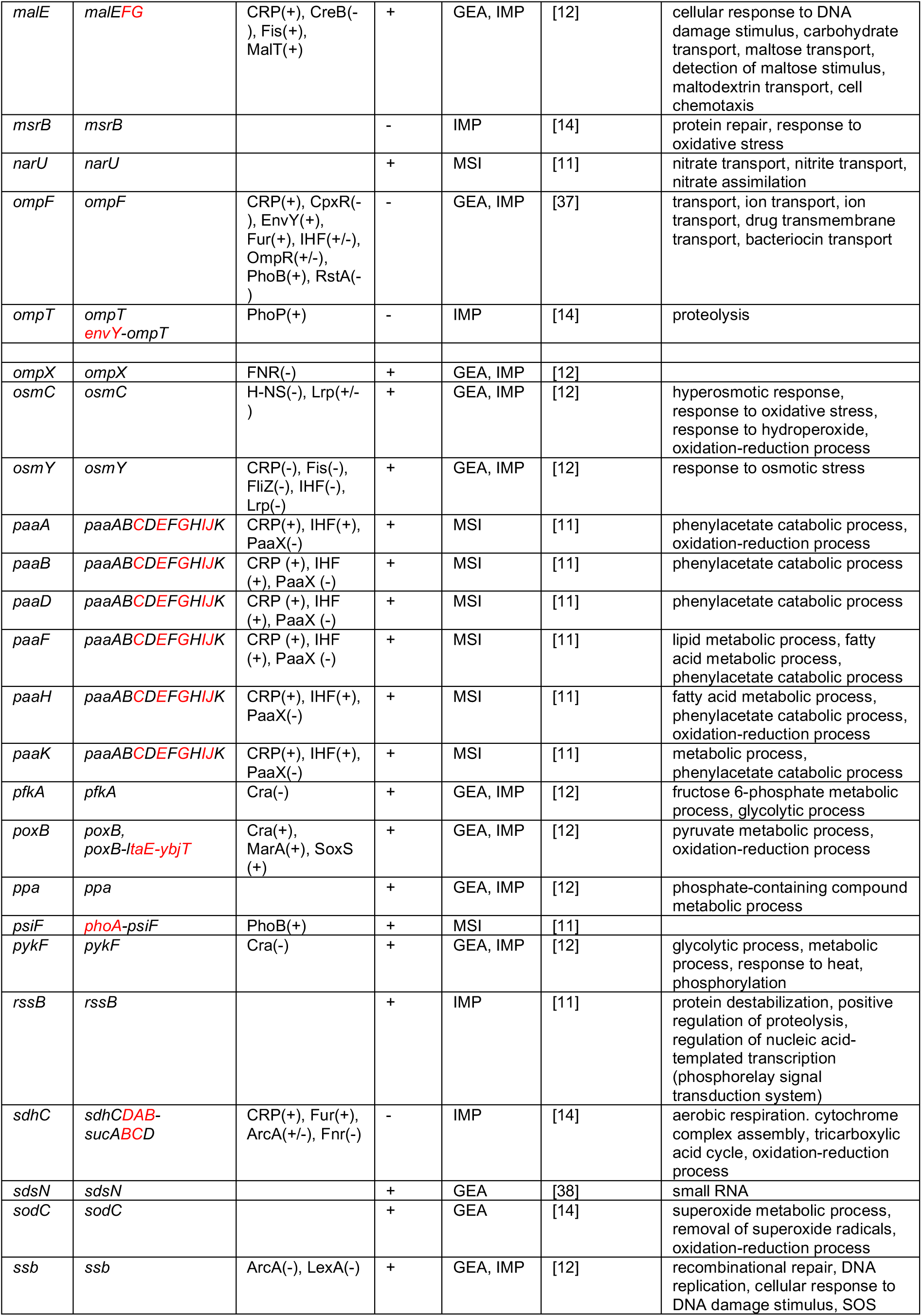
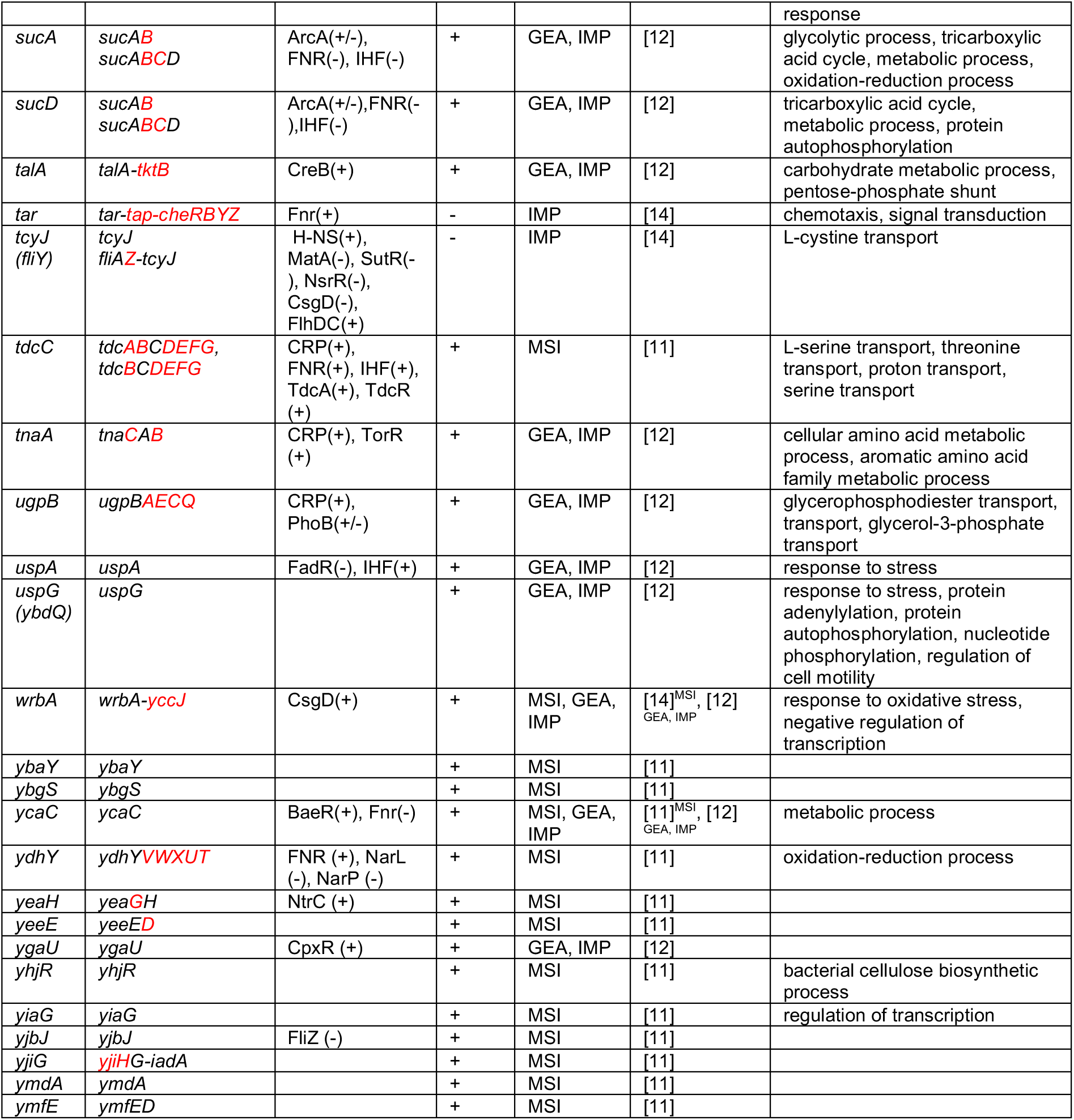
Genes regulated by Crl, TUs to which they belong (in red are possible candidates regulated by Crl, since they are controlled by Crl and σ^S^, but they did not have a change of expression in the data we evaluated), TFs regulating the TU, the effect of Crl, evidences, references, and associated GO terms. Experimental evidence types supporting regulation by Crl: APPH = assay of protein purified to homogeneity; GEA = gene expression analysis, transcriptional fusions (*lacZ*), MSI = mapping of signal intensities, such as RNA-seq or microarray analysis; IMP = inferred from mutant phenotype (such as a mutation of a TF that has a visible cell phenotype and it is inferred that the regulator might be regulating the genes responsible for the phenotype). Growth conditions were 30°C, as the stationary phase was induced for all experiments. All experiments were done with *E*. *coli* K-12 or derivative strains. All this information can be found in RegulonDB.

The regulatory network generated was displayed using the Cytoscape program, version 3.3.0 [17], with information obtained in the identified papers as well as information contained in RegulonDB [3]. Genes under Crl control were classified based on Gene Ontology (GO) annotations (http://www.geneontology.org/) using the Gene Association Format (GAF 2.0) as well as the Multifun classification scheme [18]. An enrichment analysis was carried out to find overrepresented annotations, using the PANTHER Classification system program, version 12.0. Based on our results, we selected biological processes and *E*. *coli* as parameters [19, 20]. In addition, we used KEGG to categorize the functions of GOs (http://www.genome.jp/kegg-bin/show_organism?menu_type=pathway_maps&org=eco) [21].

### Identification of Crl homologs

The Crl protein sequence of *E*. *coli* K-12 (ID: P24251) was used as the seed to scan all the bacterial and archaeal genomes via a BLASTp search [22] (E-value ≤ 10^-3^ and coverage ≥ 60%). All proteins were compared and aligned using the Muscle algorithm [23] with default parameters, and results were manually edited with the program Jalview. Finally, a phylogeny was inferred by the maximum likelihood method with 1,000 replicates by using the program MEGA [24] and the Tamura-Nei model.

### Identification of orthologous genes

Orthologous genes have been classically defined as encoding proteins in different species that evolved from a common ancestor via speciation [25] and have retained the same function. In this work, orthologs were identified by searching for bidirectional best hits (BDBHs) in other organisms [26].

### Clustering of orthologous genes

In order to evaluate the taxonomical distribution of the genes belonging to the Crl regulon, their orthologs were traced along 18 archaeal and 69 bacterial taxonomical divisions. To this end, the relative abundance of the orthologs was calculated as the fraction of genomes in the group that had one ortholog, divided by the total number of genomes per phylum, i.e., the ratio (total number of orthologs in a phylum) / (total number of organisms in phylum). Thus, the value goes from 0 (absence of orthologs) to 1, or 100%, when all organisms in the division contain an ortholog. The corresponding matrix was analyzed with a hierarchical complete linkage-clustering algorithm with correlation uncentered as the similarity measure. We used the program MeV to perform the analyses (http://www.tm4.org/mev).

## Results

### A total of 86 genes were identified as members of the Crl regulon

Available information regarding the Crl regulon was integrated through an exhaustive review of the literature. In this regard, diverse experimental evidences were considered significant for determining the association between the regulated genes and Crl protein regulator, such as gene expression analysis (transcriptional fusions), mapping of signal intensities (RNA-seq or microarray analysis), and inferences made from a mutant phenotype (mutation of a TF with a visible cell phenotype), among other analyses. In total, 52 genes of the 86 were identified in this work as new members of the σ^S^ sigmulon based on microarray data and *crl rpoS* double mutants [9-14, 27-29] (Supplementary material Table S1). From the 86 genes identified as members of this regulon (see Table 1 and Figure 1), 34 have a σ^S^-type promoter that were experimentally determined [3] and 8 genes have 13 σ^S^-type promoters that were predicted by computational approaches. These 86 genes are organized in 77 transcription units (TUs), where 52% are TUs with only one gene.

**Figure 1.**
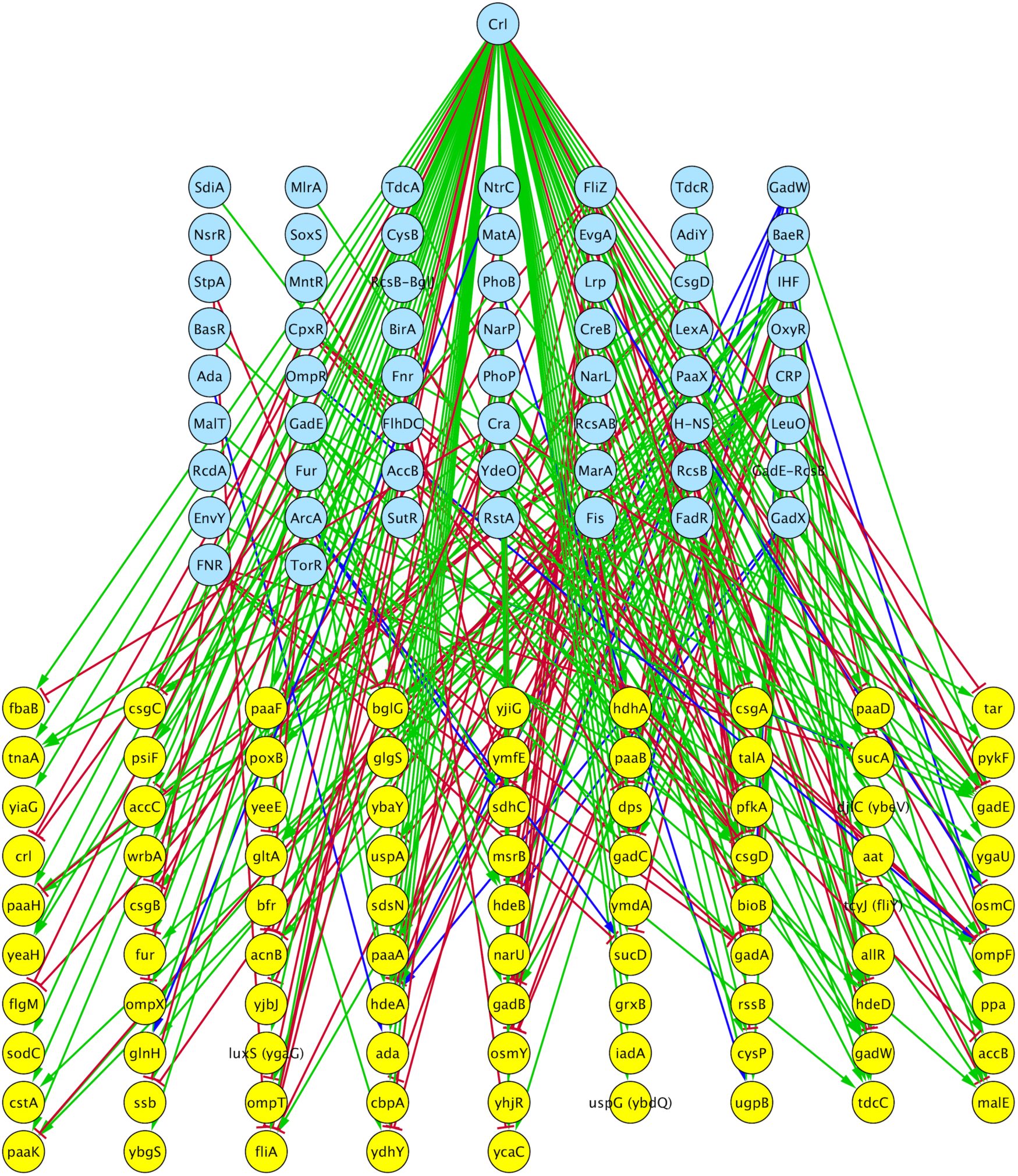
Crl regulatory network in *E*. *coli* K-12. Crl is shown in the upper part in the light blue circle. TF are shown in the middle of the network in light blue circles, and genes under Crl control are shown in yellow circles at the bottom. The effects of both Crl and TFs are shown as green solid lines for activation and red solid lines for repression.

Previously, genes under the control of Crl were classified in four main categories depending on their role(s) in the cell: DNA metabolism, central metabolism, response to environmental modifications, and miscellaneous [12]. Based on Gene Ontology (GO) annotations, multifunctional classification, and KEGG pathway maps to categorize functions, Crl-regulated genes appear to be involved in metabolic processes such as energy metabolism, amino acid, carbohydrate, and lipid metabolism, and biosynthetic processes such as glycan biosynthesis and biosynthesis of other secondary metabolites, among other metabolic processes. This function correlates with results of the enrichment analysis using PANTHER, which showed that catabolic processes, metabolic processes, and cellular responses to xenobiotic stimuli were overrepresented among the functions associated with genes under the control of Crl (See Figure 2).

**Figure 2.**
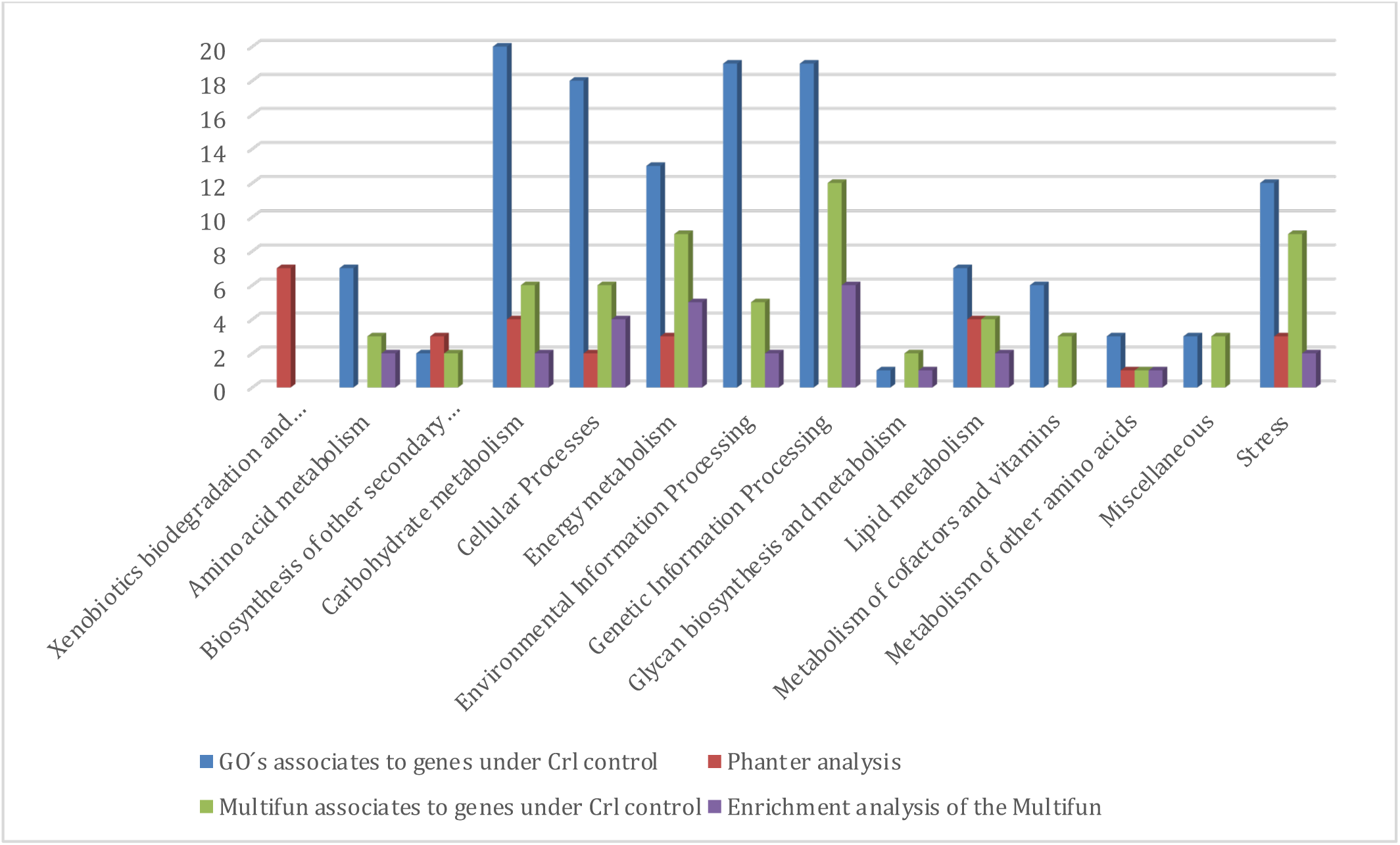
Gos and Multifun-associated genes under Crl control and enrichment analysis with the PANTHER classification system and Multifun. Categories of KEGG used to classify GOs and Multifun terms are shown on the X-axis, and the number of GOs associated with each category are shown on the Y-axis.

In general, genes under Crl control are involved in regulating many aspects of cellular metabolism through Crl’s interaction with a supset of genes of the σ^S^ regulon [9] in addition to quorum sensing playing a major role in cell-to-cell communication during stationary phase and in different processes such as biofilm formation or virulence, and also transporters [12] and genes involved in the uptake and utilization of β-glucosides [29].

### Composition of the Crl regulon

In order to determine whether additional TFs also regulate the genes under the control of Crl, RegulonDB was used to evaluate how genes associated with Crl are also regulated by alternative TFs or sigma factors. A total of 24 genes were identified as exclusively controlled by Crl, whereas 62 are regulated by TFs beyond Crl (Supplementary material Table S2). In this regard, 55 different TFs are involved in the regulation of genes associated with Crl, including Crp, IHF, H-NS, Fis, FNR, ArcA, GadX, GadW, GadE, and CsgD (Table 1), suggesting that all genes regulated by Crl are also involved in multiple functions beyond the stationary phase. It is interesting that six of seven global regulators identified in the regulatory network of *E*. *coli* are also associated with the set regulated by Crl. In addition, 19 genes of the total of Crl-regulated genes are regulated by one TF, 11 by two TFs, and 14 by three different TFs. Finally, 73 (85%) genes are regulated positively, whereas 12 (15%) genes are regulated negatively (Table 1). The predominance of positive regulation suggests that genes associated with this regulon are in high demand [30], and the activities of their proteins are enhanced to contend with varied environmental stimuli. Thirty-four of the 86 genes have a σ^S^-type promoter that was experimentally determined (RegulonDB). Finally, the promoters of 52 genes identified as members of Crl and of the σ^S^ sigmulon, based on transcriptional fusions and microarray analysis data, remain to be experimentally determined [3]. These findings suggest that Crl was probably recruited to regulate genes under previous regulation within the σ^S^ sigmulon.

### Phylogenetic analysis of Crl

In order to evaluate the phylogenetic history of Crl across the bacterial and archaeal cellular domains, its homologs were identified as described above in the Methods section, and a phylogenetic tree with maximum likelihood was generated (Figure 3). From this analysis, we found that Crl and its homologs are distributed almost exclusively among *Gammaproteobacteria* but do not share homology with proteins from other taxonomical divisions, as has been previously noted for *E*. *coli*, *Vibrio* spp., *Citrobacter* spp., *Salmonella* spp., and *Enterobacter aerogenes* [29]. Additional information suggests that Crl is less widespread and less conserved at the sequence level than σ^S^ [31]. In this regard, four conserved residues (Y22, F53, W56, and W82) are important for Crl activity and for Crl-σ^S^ interaction but not for Crl stability in *S*. Typhimurium [31]. On one hand it is probable that Crl homologs exist in some σ^S^-containing bacteria; however, some species might use alternative strategies to favor σ^S^ interaction with the core of the RNAP [31]. Therefore, our phylogenetic analysis suggests that Crl is a protein conserved and constrained to *Gammaproteobacteria*, such as in *Vibrio* spp., *Klebsiella* spp., *Enterobacter* spp, and *Escherichia* c*oli*. In addition, this TF was found in low copy numbers, i.e., one member of *crl* per genome. This information, together with the distribution of σ^S^, suggests that the regulator was recruited as an element to regulate a supset of σ^S^- regulated genes in *Gammaproteobacteria*. This result opens the question of whether genes regulated by Crl are also constrained to this taxonomical division.

**Figure 3.**
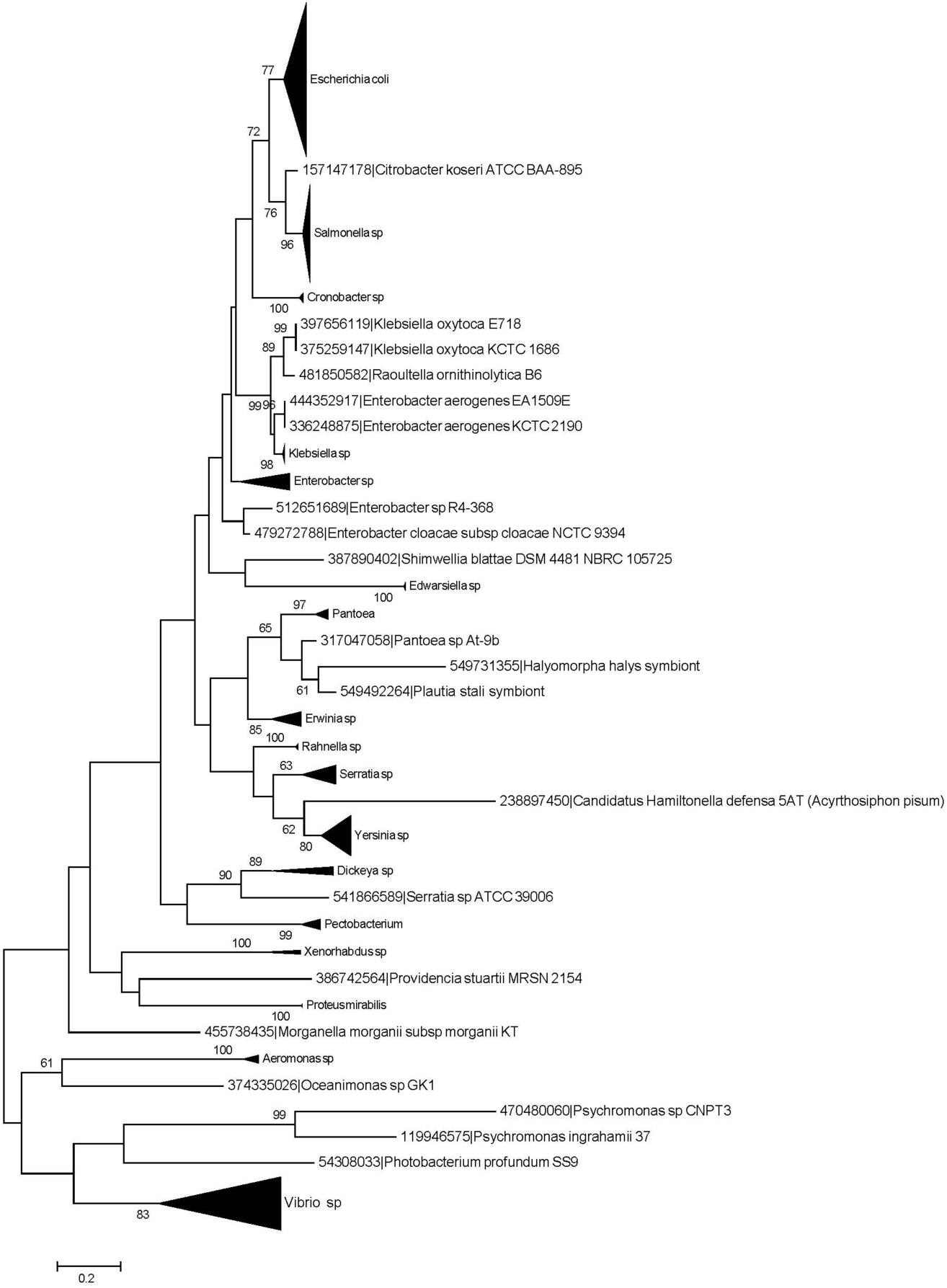
Phylogenetic tree based on Crl of *E*. *coli* and homologs in other organisms generated via maximum likelihood analysis, with 1,000 replicates. Species with bootstrap values higher than 60% are displayed. The black triangles to the right of the branches indicate multiple species for those genera.

### Taxonomical distribution of Crl-regulated genes

Based on the identification of orthologs of 86 Crl-regulated genes, we evaluated their taxonomical distribution across archaea and bacteria sequence genomes, as described in Methods (See Figure 4). Based on a taxonomical profile, we determined that the evolution of the Crl regulon seems to have involved diverse losses and gains of regulatory interactions. It is possible that large portions of the regulatory network associated with Crl evolved through extensive genetic changes during the evolution of the species studied. Indeed, we suggest three main events modeled the evolution of this regulon: (i) the recruitment of a large number of genes widely distributed among *Bacteria* and *Archea*, such as those genes involved in ancient metabolic processes such as glycolysis (*fbaB*, *pykF*, *pfkA*, and *sucA*) and those involved in the tricarboxylic acid cycle (*gltA* and *sucD*) [32]; (ii) the recruitment of genes with a distribution pattern mainly constrained to *Proteobacteria*, with some secondary losses in other organisms, such as those genes involved in response to stress and starvation (*cstA* and *hdcA*) or cell adhesion (*csgA* and *csgB*), among others; and (iii) the recruitment of Crl as a consequence of its emergence in *Enterobacteriales*. It is interesting that Crl- regulated genes are also part of the σ^S^sigmulon, where there are no essential genes [33-35]. All these elements suggest that the Crl regulon is highly flexible for phenotypic adaptation, probably as a consequence of the diverse growth environments associated with the organisms in which members of this regulatory network are present.

**Figure 4.**
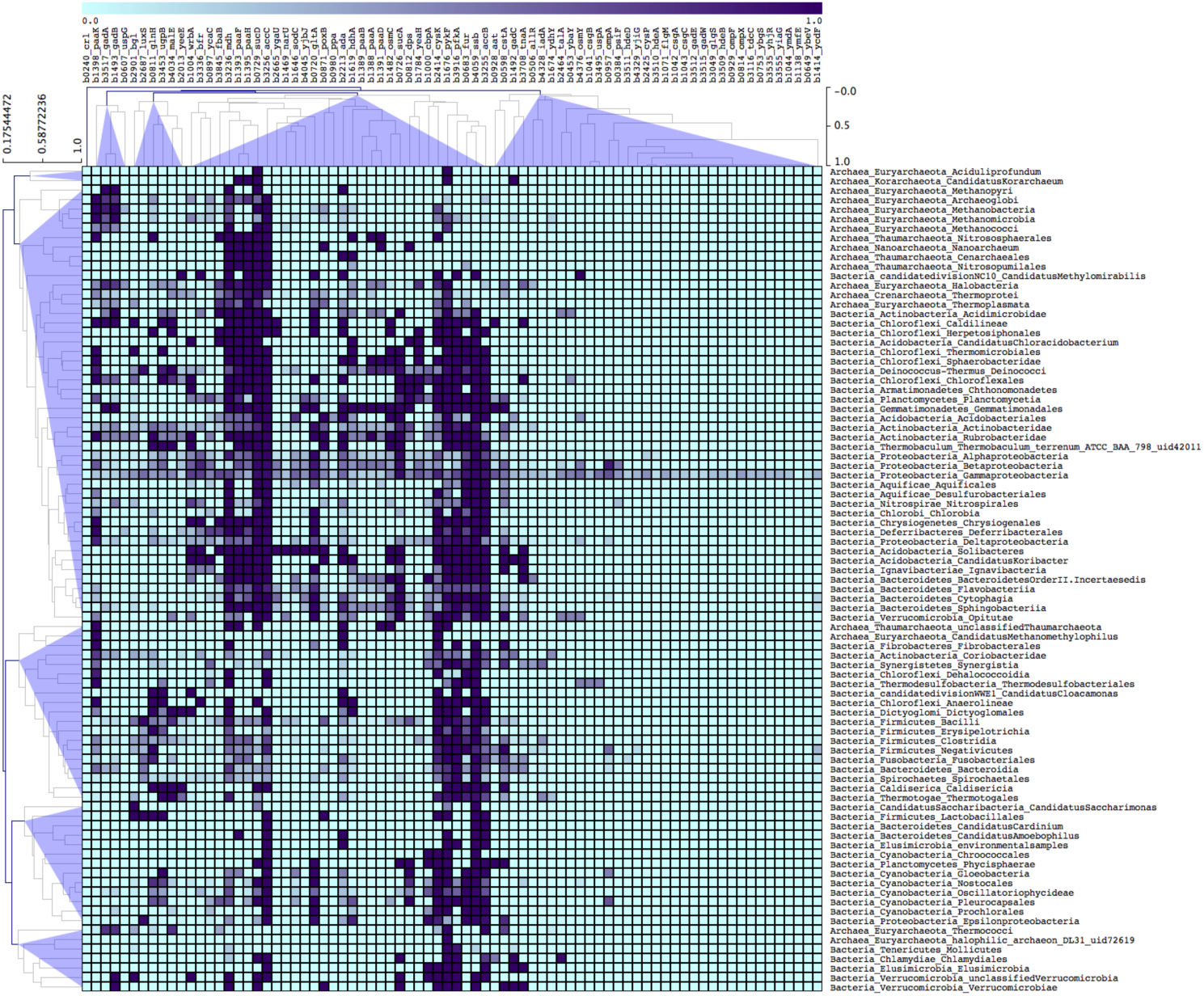
Clustering of orthologues from the perspective of *E*. *coli* K-12. A single linkage-clustering algorithm with no leaf order optimization was applied with Pearson distance as the similarity measure. The display clustering results were obtained using the MeV program (http://www.tm4.org/mev/). The conserved groups across the different taxonomic groups are indicated. Each column denotes Crl- regulated genes, whereas rows denote taxonomic groups. The bar at the top of the figure indicates the relative abundance of orthologs per group, represented as a percentage, where a value of 1 corresponds to 100% presence and 0% indicates a division without any ortholog of the Crl regulon in the taxonomic group.

## Conclusions

Crl stimulates the association of σ^S^ with the RNAP core in *E*. *coli* K-12 through direct and specific interactions, increasing the transcription rate of a supset of genes of the σ^S^ sigmulon. This TF has been described during the transition to stationary phase at low temperatures, and a recent review on the structural characterization of the Crl σ^S^ has been done [36]. In our work, based on an exhaustive literature search, we found 86 genes under the control of Crl in *E*. *coli*. These protein-coding genes were retrieved mainly from microarray and mutation analyses, among other experimentally supported evidence. These genes are associated with multiple functions, including xenobiotic processes, biofilm formation, metabolic, catabolic, and biosynthetic processes, responses to different stress conditions, and protein assembly, amino acid transport, and transcriptional processes, among others. The diverse functions regulated by Crl suggest that these genes play a fundamental role in multiple functions to respond to environmental changes, mainly those associated with stationary-phase growth at low temperatures [9]. In addition, we conducted an exhaustive analysis concerning the conservation of the regulon Crl among the Bacteria and Archaea genomes, using as a reference the knowledge gathered for *E*. *coli* K-12. From this analysis, Crl was identified in low copy numbers and constrained to the *Enterobacteriales* order, whereas the homologs of all regulated genes were found to be widely distributed beyond enterobacteria, suggesting that Crl was recruited in a secondary event to regulate a specific supset of genes for which the stimulation of Crl and σ^S^ is necessary.

